# CRISPys: Optimal sgRNA design for editing multiple members of a gene family using the CRISPR system

**DOI:** 10.1101/221341

**Authors:** Gal Hyams, Shiran Abadi, Adi Avni, Eran Halperin, Eilon Shani, Itay Mayrose

## Abstract

The discovery and development of the CRISPR-Cas9 system in the past few years has made eukaryotic genome editing, and specifically gene knockout for reverse genetics, a simpler, efficient, and effective task. The system is directed to the genomic target site by a programmed single-guide RNA (sgRNA) that base-pairs with the DNA target, subsequently leading to site-specific double-strand breaks. However, many gene families in eukaryotic genomes exhibit partially overlapping functions and, thus, the knockout of one gene might be concealed by the function of the other. In such cases, the reduced specificity of the CRISPR-Cas9 system, which may lead to the cleavage of genomic sites that are not identical to the sgRNA, can be harnessed for the simultaneous knockout of multiple homologous genes. Here, we introduce CRISPys, an algorithm for the optimal design of sgRNAs that would potentially target multiple members of a given gene family. CRISPys first clusters all the potential targets in the input sequences into a hierarchical tree structure that specifies the similarity among them. Then, sgRNAs are proposed in the internal nodes of the tree by embedding mismatches where needed, such that the cleavage efficiencies of the induced targets are maximized. We suggest several approaches for designing the optimal individual sgRNA, and an approach that provides a set of sgRNAs that also accounts for the homologous relationships among gene-family members. We further show by in-silico examination over all gene families in the *Solanum lycopersicum* genome that our suggested approach outperforms simpler alignment-based techniques.

**Graphical abstract:** 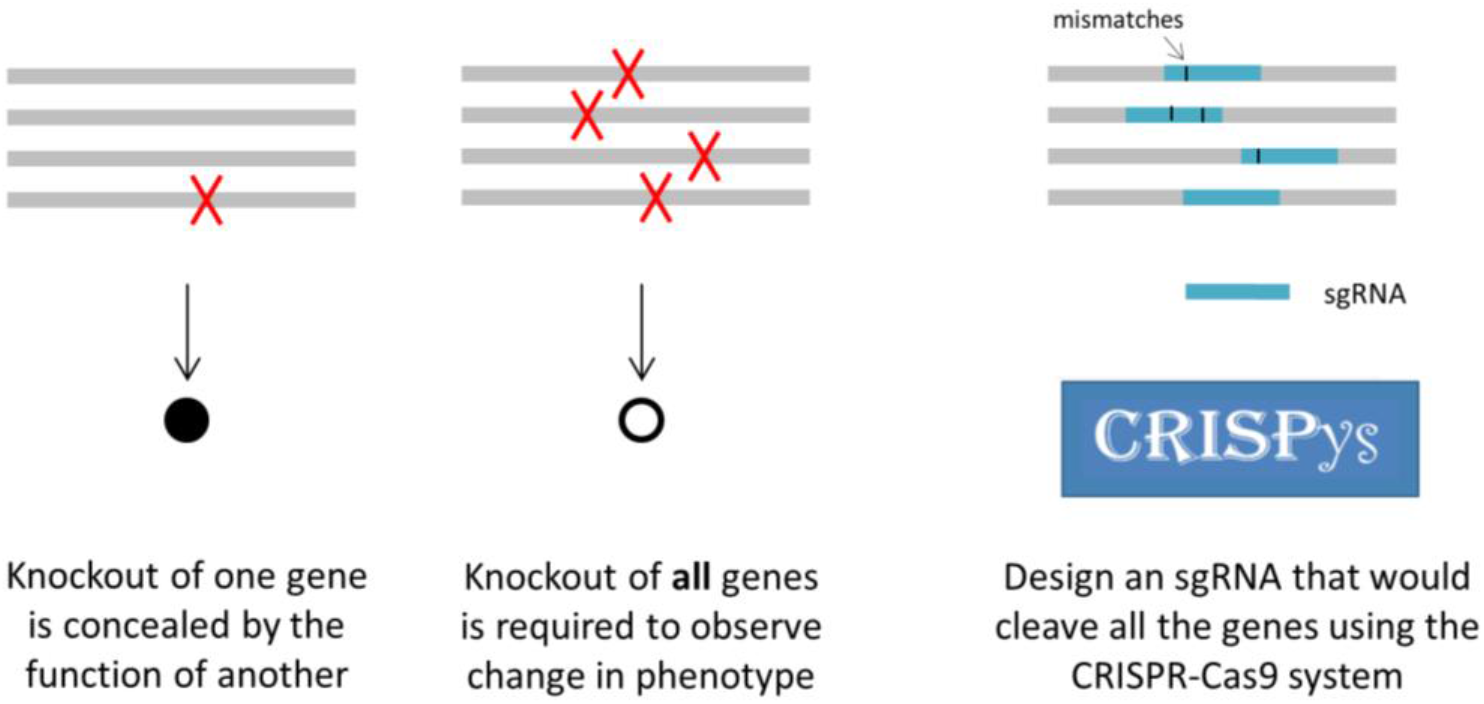

**Highlights:**

- Many genes in eukaryotic genomes exhibit partially overlapping functions. This imposes difficulties on reverse-genetics, as the knockout of one gene might be concealed by the function of the other.
- We present CRISPys, a graph-based algorithm for the optimal design of CRISPR systems given a set of redundant genes.
- CRISPys harnesses the lack of specificity of the CRISPR-Cas9 genome editing technique, providing researchers the ability to simultaneously mutate multiple genes.
- We show that CRISPys outperforms existing approaches that are based on simple alignment of the input gene family.

## Introduction

Due to extensive history of local and large-scale genomic duplications, many eukaryotic genomes harbor homologous gene families of partially overlapping functions [1–4]. For example, 72% of the protein coding genes in the Plaza 3.0 Monocots database [5], that presently covers 16 fully sequenced plant genomes, belong to paralogous gene families with at least two members. This redundancy often leads to mutational robustness such that the inactivation of one gene often results in no or minimal phenotypic consequence [2,4,6–8]. As there are no observable phenotypes for many single-gene loss-of-function mutants, it is often necessary to mutate multiple members of a gene family to uncover phenotypic consequences and to enable in-depth molecular characterization of their function. Here, we present a computational methodology that will facilitate such endeavors via genome editing techniques.

The CRISPR (Clustered Regularly Interspaced Short Palindromic Repeats) and the associated protein 9 nuclease (Cas9) system have been recently adopted as a genome editing technique of eukaryotic genomes. The target genomic DNA sequence consists of 20 nucleotides followed by the Protospacer Adjacent Motif (PAM), which is usually in the form of NGG (where N stands for any nucleotide and G for guanine). The system is directed to the genomic site using a programmed single-guide RNA (sgRNA) that base-pairs with the DNA target, subsequently leading to a site-specific double strand break four to five nucleotides upstream of the PAM sequence. This break can be repaired through the Non-Homologous End Joining (NHEJ) repair pathway, frequently resulting in a frameshift in the encoded protein, thus leading to its inactivation. Using the CRISPR-Cas9 system, DNA sequences within the endogenous genome are now regularly edited in diverse organisms as human [9], mouse [10], zebra fish [11], yeast [12], and plants [13,14], and the system is rapidly becoming the technology of choice for generating single gene knockouts for reverse genetics studies [13,15–18]. Importantly, the binding affinity of the CRISPR-Cas9 system does not require perfect matching between the sgRNA and the DNA target. Thus, in addition to cleaving the desired on-target, cleavage may occur at multiple unintended genomic sites (termed off-targets) that are similar, up to a certain degree, to the sgRNA. Several studies have demonstrated that four or more mismatches can be tolerated between the sgRNA and the cleavage site, depending on the location of the mismatches and their spatial distribution [19–21]. Indeed, it is well acknowledged that mismatches at PAM-distal positions are better tolerated than those occurring at PAM-proximal sites [20,22,23]. Thus, when designing an sgRNA for editing a single gene, two topics should be considered: the sgRNA sensitivity (maximizing the cleavage probability of the on-target) and its specificity (minimizing the cleavage probabilities of off-target sites). Several computational tools have been constructed to deal with these challenges [24–30].

To date, much effort has been devoted to refine the specificity of the CRISPR system as a means to decrease the off-target effect [31–33]. Yet, the low specificity of the system could, in fact, be harnessed to enable a rational design of an sgRNA that would simultaneously target multiple genes. An example of such a possibility was recently shown in rice, where a single sgRNA led to the modification of three homologous genes from the cyclin dependent kinase protein family [34]. The sgRNA used in that study was designed to perfectly match one of the family members, while the other two homologues possessed one and two mismatches and were silenced as a byproduct. A fourth homologue, with three mismatches, was not affected by this transfection. Yet, it is possible that considering all family members in the design process would enable the cleavage of a larger fraction of the homologous gene family and would enhance the balance between their cleavage frequencies.

One approach for accomplishing this task is to align the sequences of the given genes and then locate highly similar CRISPR-Cas9 target sites in the consensus sequence, while allowing for few mismatches between the consensus and each of the aligned sequences. This strategy is implemented in the MultiTargeter tool [35], which also accounts for CRISPR-specific considerations such as allowing mismatches only within PAM-distal nucleotides. In case that multiple potential sgRNAs are found, these are ranked according to their efficacy, as predicted by the CFD score [25]. While the MultiTargeter algorithm is very efficient in terms of computational running time, it may miss a large number of valid candidates. For example, similar 20-nt long subsequences that appear in two homologous genes but do not overlap in the resulting alignment, or that are in opposite strands, will be ignored.

Here, we present a novel method, termed CRISPys, aimed for the design of an optimal set of sgRNAs for silencing multiple members of a gene family using the CRISPR-Cas9 system. CRISPys detects highly similar sequences among the set of all potential CRISPR-Cas9 targets (i.e., sequences that are followed by a PAM site) located within the genes of interest and then designs sgRNAs that would cleave the gene set with highest efficacy. CRISPys can further incorporates any scoring function specifying the cleavage propensity of a genomic site by a given sgRNA, thus allowing flexible use of the method with the accumulation of knowledge regarding this emerging genome engineering technique. We present the utility of CRISPys by applying it in a genome-wide manner to numerous gene families in the *Solanum (S.) lycopersicum* genome and compare its performance to the existing alignment-based approach.

## Results

### 1. Algorithm description

Given a set of genes, *G*={*g_i_*}, potentially belonging to a homologous gene family, and a scoring function *φ*, we would like to identify suitable sgRNA candidates that are likely to cleave the largest number of genes in *G*. The input scoring function *φ*(*sgRNA, target*) → [0,1] specifies the estimated cleavage propensity of a genomic target by a given sgRNA, such as the CFD score [25], CROP-IT [36], Optimized CRISPR Design [27], or CRISTA [37].

#### 1.1 Targets tree construction

Ideally, we would like to examine all possible sgRNA candidates and identify the one that would cleave the given gene set with highest propensity. This, however, entails the examination of an exceedingly large number of sgRNA possibilities (4^*l*^, where *l* is the length of the sgRNA, typically *l*=20) leading to computationally intractable running time. The examined set is thus narrowed to the sgRNAs that are potentially relevant to the input gene set. To this end, all potential targets within the genes in *G* are clustered into a hierarchical tree structure that specifies the similarity among the targets as follows. First, for each gene *g*_*i* ∈ *G*, all potential targets are extracted. By default, these are defined as 20-nt long sequences upstream to an NGG motif; additional PAM motifs (e.g., NAG) or other sequence lengths (17-23) can be specified. Second, the entire set of potential targets, *T*, are clustered using the UPGMA hierarchical clustering algorithm [38] and placed in a tree structure, such that more similar targets (represented by the tips of the tree) are placed closer to each other on the tree. The input pairwise distance matrix for the UPGMA clustering is computed using the scoring function *φ*(*sgRNA, target*) that is transformed into a distance metric (see Methods). An internal node, *a*, in this tree induces a set, *T^a^*, of potential targets that are the descendants of this node. As *a* gets closer to the root, the size of *T^a^* increases such that the contained targets become less similar and are less likely to be cleaved by a single sgRNA. We also denote by *G^a^* the group of genes to which the targets in *T^a^* belong. Figure 1 presents an example of a target tree for a family of five genes, each represented by a single potential target.

**Figure 1.**
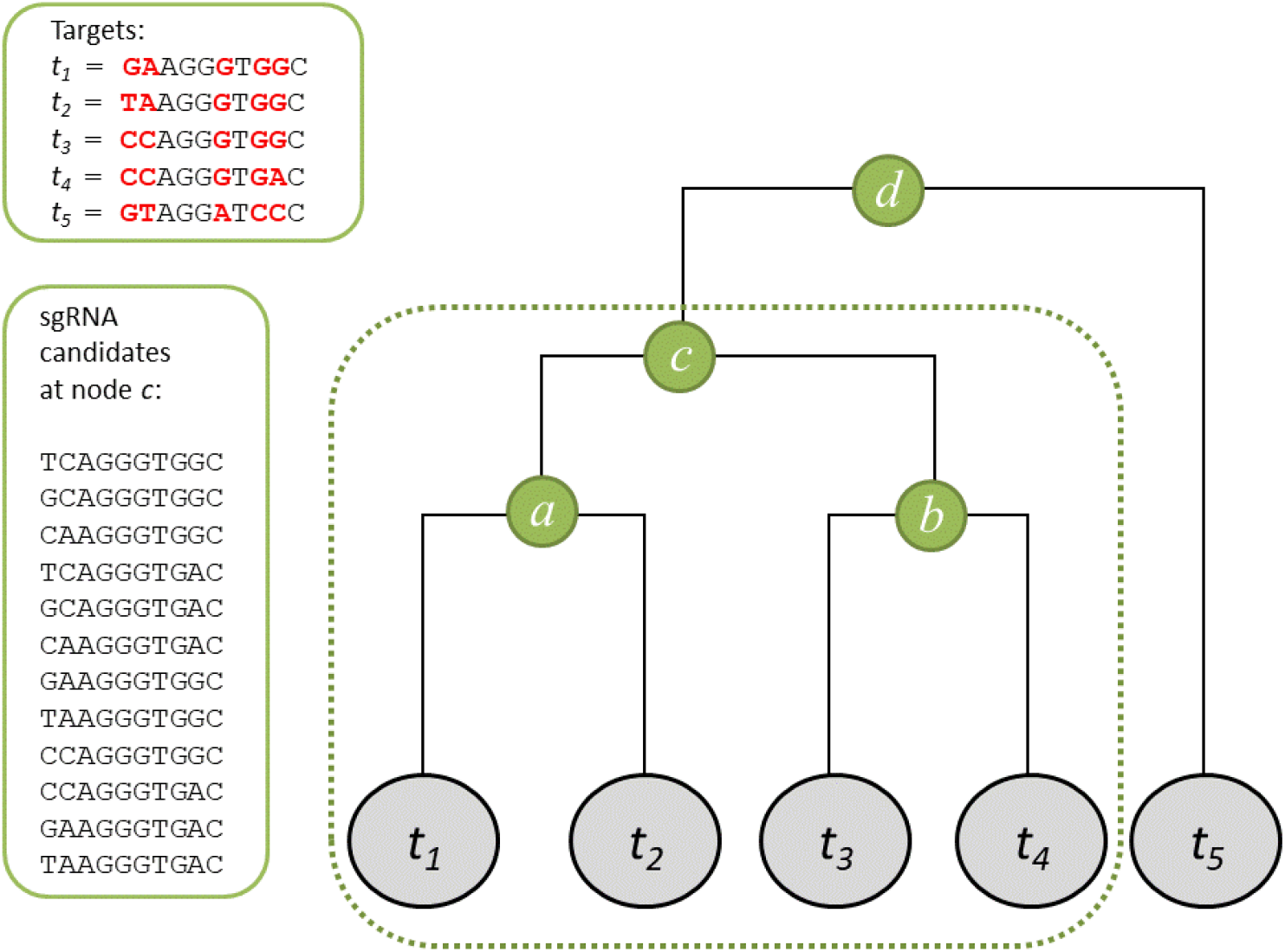
An illustrative example of a target tree used for sgRNA design. A group of five genomic targets (*t_1_-t_5_*) is hierarchically clustered according to pairwise distances. Each gene is represented by a single target and only the first 10 positions upstream of the PAM site are shown for each target (top-left panel). Nucleotides at polymorphic positions are indicated in red. The target set induced by internal node *a, T^a^*, includes *t_1_* and *t_2_* while *T^d^* includes all five targets. The numbers of polymorphic sites within *T^a^, T^b^*, and *T^c^* are 1, 1, and 3 and are below the *k* cutoff (set here to be 4), while the number of polymorphic sites within *T^d^* is above *k*. Possible sgRNA candidates are considered based on all possible combinations for the polymorphic sites in *T^c^*, as listed at the panel to the left of the tree.

#### 1.2 Target-tree traversal and identification of sgRNA candidates

Once the targets tree is constructed, the algorithm proceeds by traversing the tree in a post-order manner, identifying sets of targets for which sgRNA candidates will be designed. Specifically, upon reaching an internal tree node *a*, the number of polymorphic sites in the induced target set, *T^a^*, is calculated. If this number is above a cutoff *k*, the search does not proceed up the tree and potential sgRNAs are designed based on each of the descendent subtrees. Otherwise, tree traversal is continued. In all our analysis, *k* was set to 12 since *k* ≥ 13 led to exceedingly long running time without producing any improvement in the assessed efficacies of the additional sgRNA candidates.

For each target set identified, sgRNA candidates are designed by enumerating over all possible combinations of the polymorphic sites found within it (Figure 1). The efficacy of each candidate sgRNA s to target the genes in *G* is then assessed. Specifically, *φ*_*s* (*g*_*i*), the cleavage propensity of gene *g*_*i* ∈ *G* by sgRNA *s* is computed by considering all possible targets that belong to *g*_*i* as follows:

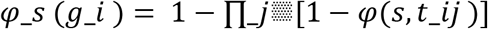

where *φ*(*s, t*_*ij*) is the cleavage propensity of the *j*’th target site of gene *g*_*i* by sgRNA *s* (as calculated by the input scoring function). This way, as the number of similar targets in *g*_*i* increases so does the propensity that the gene is cleaved by the specified sgRNA; in case a gene has only one target, *φ*_*s* (*g*_*i*) reduces to *φ*(*s, t*_*i*1).

Next, we compute Φ_s (*G*); the efficacy of each sgRNA, *s*, to cleave the set of genes; and choose the best candidate. Intuitively, we would have liked to compute the joint propensity of the given sgRNA to cleave the entire set of genes, which translates into the following function:

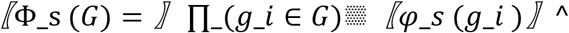

Yet, the propensity to cleave all genes dramatically decreases when no sgRNA is decently suitable for all of the members. In such cases, it becomes irrelevant to use this function as an optimality criterion among the sgRNAs candidates. Thus, we implemented two alternative criteria for computing Φ_s (*G*):

*Criterion 1:* for each sgRNA *s*, compute the cleavage expectation across **all** genes in *G*: 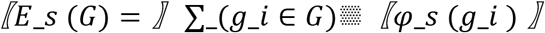. The optimal sgRNA candidate is defined as

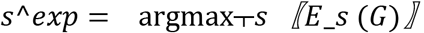

*Criterion 2:* instead of optimizing the cleavage of all genes, concentrate on those with high cleavage propensity. This criterion thus ignores genes whose propensity of cleavage by a given sgRNA is below a certain threshold. Specifically, we define *G*_*s*^*Ω* as the group of those genes that are expected to be cleaved by sgRNA s above a given threshold *Ω*:

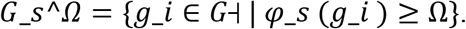

Let *c*_*s*^*Ω* be the size of this group: *c*_*s*^*Ω* = |*G*_*s*^*Ω*| and let *c*_*max*^*Ω* be the highest *c*_*s*^*Ω* value computed for the group of input genes G. Because there may be multiple sgRNAs with this *c*_*max*^*Ω* value, the optimal sgRNA is chosen as the one with the highest propensity to cleave all *c*_*max*^*Ω* genes:

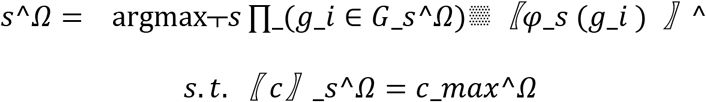

Notably, the use of this criterion necessitates the use of an arbitrary threshold (*Ω*). Setting *Ω* to 0.0 results in all input genes affecting the design, while setting *Ω* = 1 practically seeks the sgRNA that perfectly matches the largest number of genes. To set *Ω* to realistic values, the thresholds used for performance evaluation (see section below) were determined according to an experimental dataset that was profiled by the genome-wide detection technique, GUIDE-Seq [19]. Specifically, the data in that study are composed of a collection of 10 sgRNAs that overall cleaved 413 targets throughout the human genome. For each of these validated targets and its corresponding sgRNA, we calculated the predicted cleavage propensity using the CFD scoring function [25]. The upper 10^th^ (Ω = 0.66) and 50^th^ (Ω = 0.43) percentiles were then chosen as the stringent and permissive thresholds, respectively.

### 2. Number of genes predicted to be cleaved by the best sgRNA

To demonstrate the utility of CRISPys and to evaluate its different design strategies *s*^*exp* and *s*^*Ω*, we applied it to all 3,697 gene families of size 2-10 within the tomato (*Solanum lycopersicum*) genome. The classification of genes to families was taken from the Plaza plant comparative genomics database [5]. Using the *s*^*exp* design—that considers the entire gene set—the expected number of genes predicted to be cleaved increases with the number of genes that are included in the family (Figure 2). The average cleavage expectation approaches a plateau, such that the expected number of cleaved genes by the optimal sgRNA remains around 4 for families of size eight or higher. A similar trend is obtained using the *s*^*Ω* design (Figure 3), which considers only genes whose propensity of cleavage is high. As expected, as the family size increases, the optimal sgRNA is predicted to cleave more genes, although the tendency to cleave *all* genes in the family decreases. For example, in 87% of the families of size two, the best sgRNA could cleave all family members. This percentage decreases to 10% for families of size six, while for all 50 gene families of size ten no such sgRNA could be obtained. Finally, an asymmetric tradeoff between the two design strategies is demonstrated in Table 1. When all genes in the family are considered, the cleavage expectation of the *s*^*exp* candidate is higher compared to that of *s*^*Ω* (2^nd^ and 4^th^ columns). Yet, when only genes that are predicted to be cleaved with high propensity are considered, the *s*^*exp* candidate would cleave fewer genes than the *s*^*Ω* (3^rd^ and 5^th^ columns).

**Figure 2.**
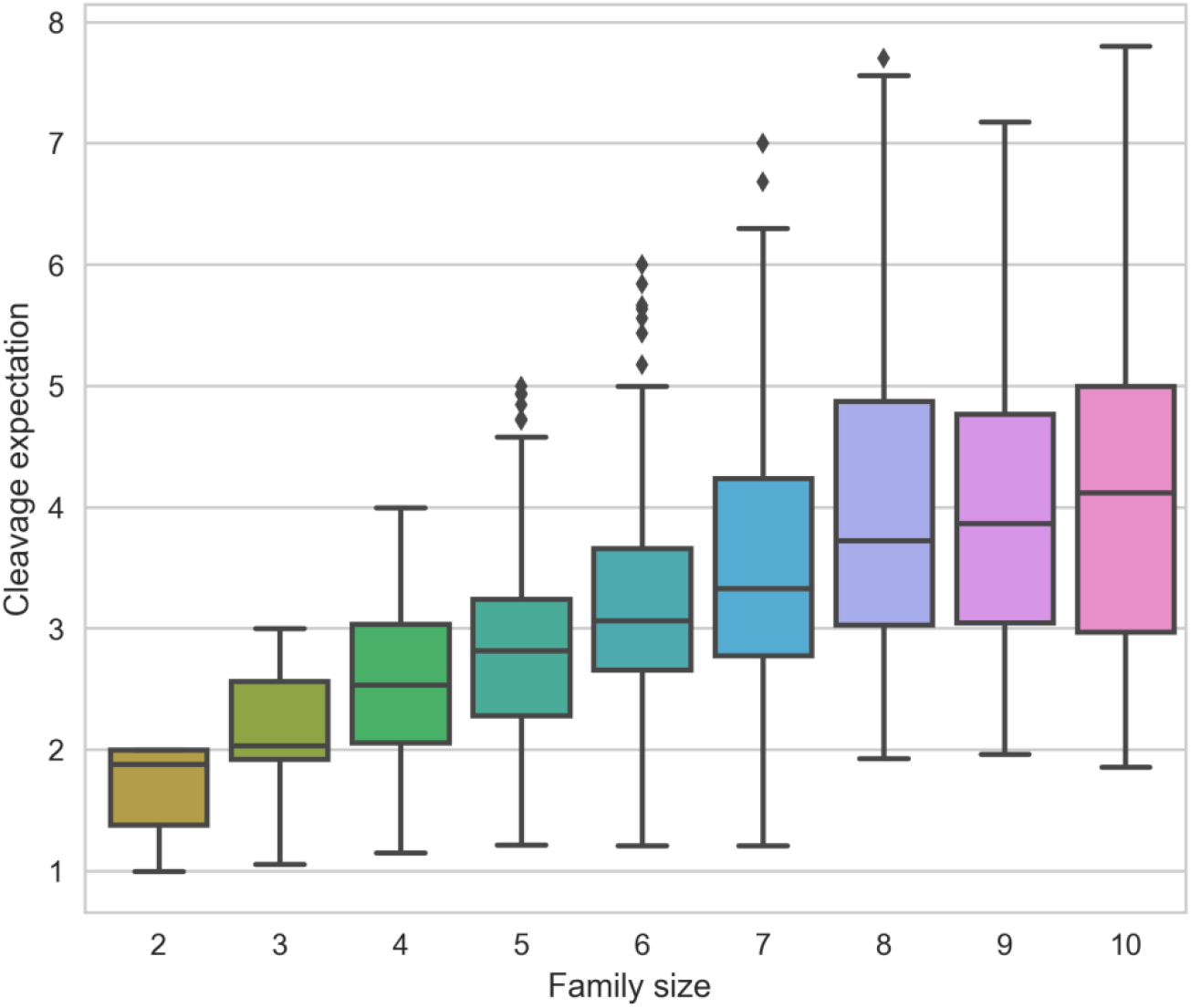
CRISPys results across the genome of *S. lycopersicum* using the ***s*^*exp*** design strategy. A box plot describing the expected number of cleaved genes by the *s*^*exp* candidate, over different family sizes. The lines within the boxplot represent the 1^st^, 2^nd^, and 3^rd^ quartiles and diamonds above the upper line represent outliers.

**Figure 3.**
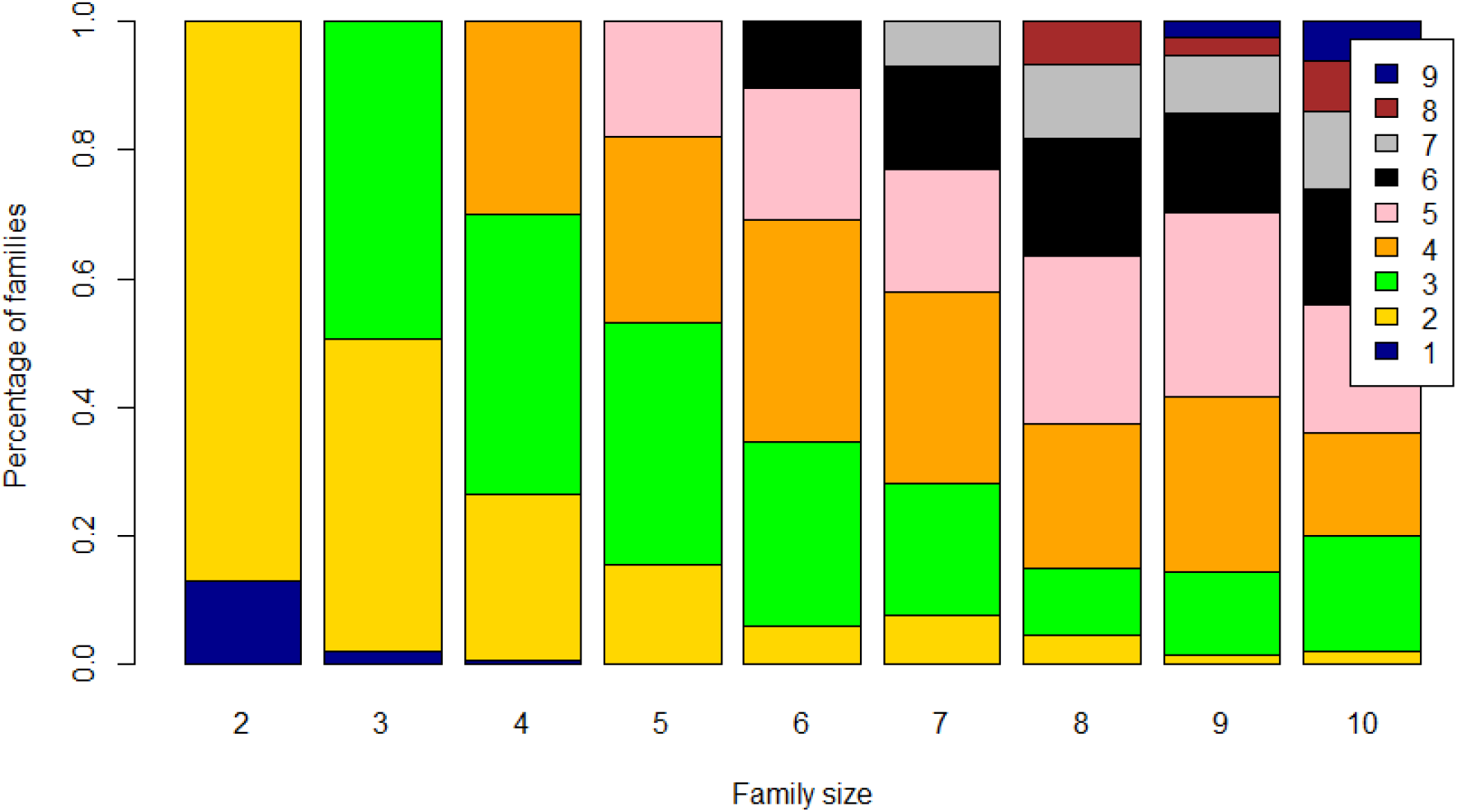
CRISPys results across the genome of *S. lycopersicum* using the ***s*^*Ω*** design strategy. The number of genes predicted to be cleaved for all gene families of size 2-10. Each bar represents the results obtained for a given family size. The color bar at the right of the panel specifies the number of genes predicted to be cleaved by the *s*^*Ω* candidate. These results were computed with threshold *Ω* = 0.43.

**Table 1.**
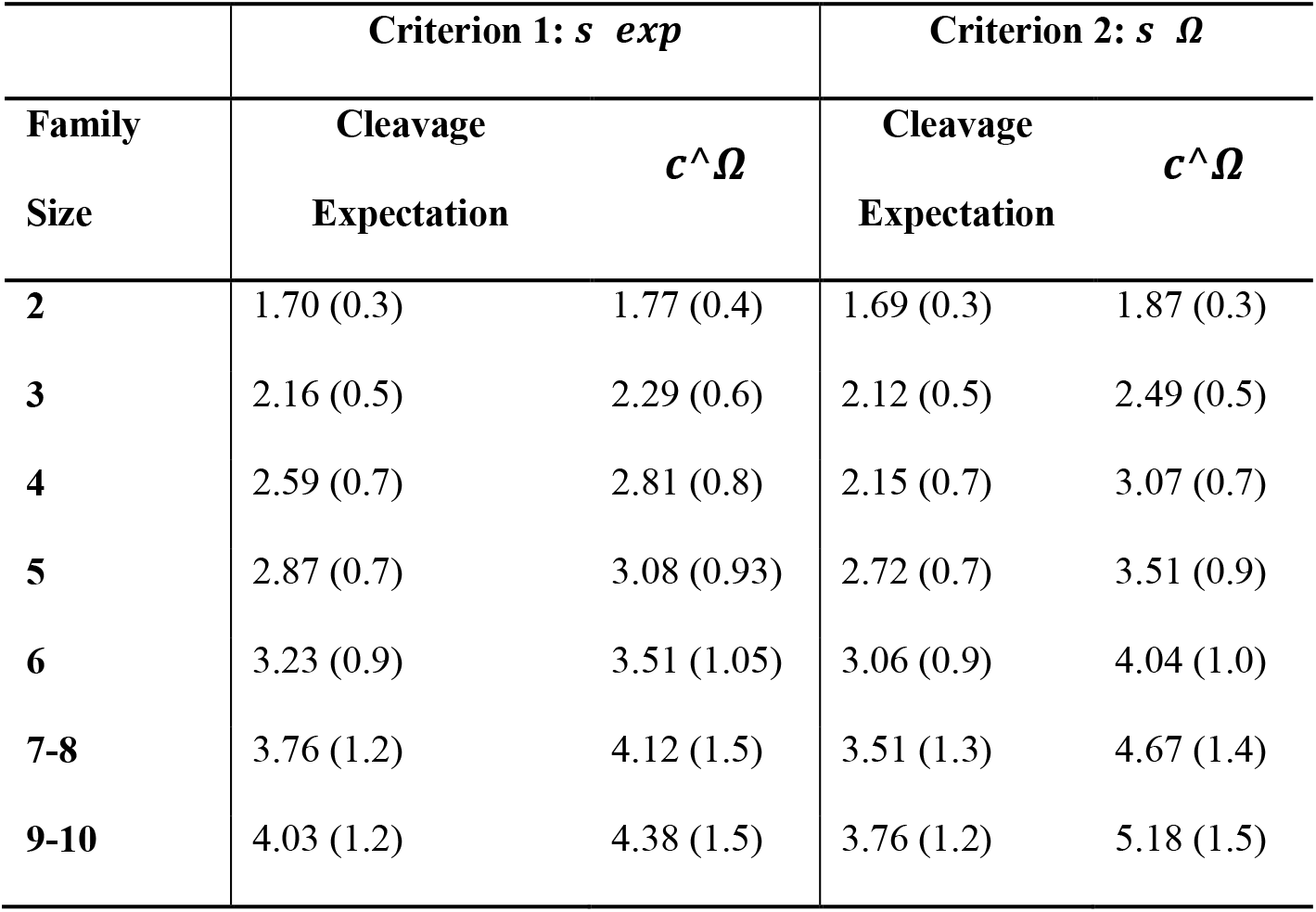
Comparison between the decision criteria of CRISPys. For each family size of 2-10 genes in the genome of *S. lycopersicum*, CRISPys was applied with either the *s*^*exp* or *s*^*Ω* criterion. Cleavage expectation is the average expected number of genes predicted to be cleave across all families of that size and *c*^*Ω* is the average number of genes that are predicted to be cleaved above a threshold of *Ω* = 0.43 by each design strategy. The standard deviation of each statistic is given in parentheses. See Supplementary Table S1 for results obtained using other *Ω* values.

### 3. Comparison between CRISPys and a consensus-based approach

To compare the results obtained using CRISPys to a consensus-based approach, we applied the MultiTargeter tool [35] to the same set of families within the *S. lycopersicum* genome. Notably, MultiTargeter identifies a promising sgRNA only when all family members are predicted to be cleaved by a single sgRNA while no results are obtained otherwise. Thus, in this comparison we defined the prediction of CRISPys as successful if the predicted cleavage propensity of each of the genes by the designed sgRNA is above a threshold of *Ω* = 0.43 (as was determined based on experimental evidence; see section 1.2). This definition was applied to both the *s*^*exp* and *s*^*Ω* design criteria of CRISPys. In comparison, MultiTargeter allows one mismatch in the eight positions furthest of the PAM, which translates to a more lenient lower bound of *Ω* = 0.35 using the CFD scoring function [25]. This analysis revealed several important observations (Table 2). First, CRISPys produced a successful prediction for all gene families for which MultiTargeter returned a result. Second, for all gene families considered, CRISPys obtained a larger fraction of successful predictions. Third, the percentage of successful predictions by all alternatives decreases as the family size increases. Yet, this decline is much shallower for the two design criteria of CRISPys, particularly *s*^*Ω*, as compared with that obtained using MultiTargeter. For example, for families with two genes, the ratio between the number of successful predictions between CRISPys *s*^*Ω* design and MultiTargeter is 2.49, while this ratio arises to 7.75 and 10.0 for families of size four and six, respectively.

**Table 2.** Comparison between CRISPys and MultiTargeter. Number of families in the genome of *S. lycopersicum* for which an sgRNA that can cleave all family members could be successfully designed using MultiTargeter and the two decision criteria of CRISPys. For MultiTargeter, percentage of families in each size signifies the fraction of runs for which a result could be obtained. For CRISPys, a prediction is considered successful if the cleavage propensity of all genes is above *Ω* = 0.43.

| Family size | #Families | CRISPys | CRISPys $s \ \Omega$ | MultiTargeter |
| --- | --- | --- | --- | --- |
|  |  | *s exp* |  |  |
| 2 | 1690 | 77% | 87% | 35% |
| 3 | 757 | 36% | 51% | 10% |
| 4 | 432 | 22% | 31% | 4% |
| 5 | 290 | 9% | 18% | 1% |
| 6 | 182 | 4% | 10% | 0.5% |
| 7-8 | 219 | 4% | 7% | 0.5% |
| 9-10 | 127 | 1% | 1% | 0% |

## Discussion

In this work, we presented CRISPys, a novel computational method that utilizes the nonspecificity of the CRISPR-Cas9 system for the design of an optimal sgRNA that would most efficiently mutate multiple members of a gene family. The efficiency is computed by one of several scoring functions regarding the cleavage propensity of genomic sites by a given sgRNA. This allows the user to prioritize the considerations of choice. For example, the function provided by the ‘Optimized CRISPR Design’ [27] is practically dichotomous, as it assigns a score of 1.0 to all on-targets and to some targets with a single mismatch, while the score of nearly all other targets approaches zero. Using this function will thus consider for each candidate sgRNA only the most definite targets. The CFD score [25], on the other hand, is more delicate and is sensitive to the type and position of each mismatch, and its use will thus consider a broader collection of potential targets. Aside from functions that consider the pairwise similarity between the sgRNA and the DNA target, CRISPys can also integrate functions that account for additional genomic features (e.g., the GC content or the DNA rigidity surrounding the target site [39]. Notably, using one scoring function yields different results than using another, and as the research of the CRISPR-Cas9 system evolves, any new function can be easily incorporated within CRISPys. This flexibility in the underlying scoring function further enables CRISPys to readily design sequences for guiding any of the emerging CRISPR nucleases variants, including the recently studied class-2 Cas proteins [40–42].

The sgRNA design by CRISPys is not only dependent on the scoring function, but also on the criterion by which one chooses to select the optimal sgRNA. Two alternative strategies were presented. A user that is interested to maximize the number of family members to be cleaved should, in principle, select the sgRNA with the highest cleavage expectation over the entire set of input genes, as computed using the *s*^*exp* strategy. Yet, at least at present, the large gaps of knowledge surrounding the CRISPR-Cas9 mode of action and the noisy experimental procedures that are used for its evaluation translate to scoring functions with large degrees of uncertainty. Given the costly (and timely) experimental resources that are needed to validate a successful cleavage, researchers are most often interested in focusing their efforts in validating those targets whose probability of cleavage is high. This is implemented using the *s*^*Ω* design, but necessitates the use of a pre-specified cleavage-propensity threshold above which targets are considered.

An alternative approach for targeting multiple genomic sites using a single sgRNA has been previously implemented in the MultiTargeter webserver [35]. This approach detects potential targets in the consensus sequence of a multiple sequence alignment of the input sequences. This procedure entails several difficulties. First, it relies on the ability to correctly align the input gene sequences, a procedure that is known to be error-prone [43–46]. Second, this approach entails that only sites that are aligned in the same positions are considered, and that a single distant sequence suffices to prevent the design of a proper sgRNA, even if the rest are highly conserved. Third, the consensus would assign the most abundant character at each position, while a more balanced design would disperse mismatches over the input genes, accounting for the specific penalty of each assignment. In contrast, since CRISPys first clusters all potential targets according to sequence similarity, it is not dependent on their locations or their orientation. Moreover, by incorporating any specified scoring function, CRISPys allows for a more sensitive consideration of each site. Indeed, as shown by our analysis of the tomato genome, CRISPys succeeded in providing promising sgRNA candidates for a larger number of gene families compared to the MultiTargeter consensus-based approach.

Evidently, CRISPys aims at optimizing the cleavage of the given genes by increasing the cleavage scores. However, a designed sgRNA may also cleave additional “off-target” sites, leading to the knockout of undesired genes. Therefore, sgRNA design should also minimize any off-target effects. Ideally, off-target considerations could be integrated within the computations performed by CRISPys. One option would be to balance between efficiency (maximizing cleave propensity of the input genes) and specificity (minimal cleavage of off-target sites) using a tunable parameter. This, however, requires the detection of potential off-targets for each and every sgRNA considered throughout the course of the algorithm, rendering it computationally infeasible. Alternatively, and as implemented in the online webserver, CRISPys generates a list of sgRNAs ranked according to their computed cleavage propensity. For each of these, off-target detection could be performed through a number of existing applications [26,27,30,47], thereby allowing researchers to choose the sgRNA that is most suitable for their need.

Recently, multiplex genome editing has been introduced, thereby allowing the application of multiple sgRNAs within a single construct [9,22,23,48–51]. These systems have been shown to be useful for the simultaneous knockout of multiple protein coding genes and for the deletion of noncoding RNA regions and other genetic elements [34,52]. Multiplex genome editing could be combined with the CRISPys algorithm to design a set of sgRNAs that would collectively mutate a large fraction of the input gene set. The *s*^*Ω* design option of CRISPys is particularly appealing in this regard since the *Ω* threshold could be tuned in such a way to allow a more strict (or lenient) design of sgRNAs such that each sgRNA would target a narrower (or broader) fraction of the genes. A refined tuning of the *Ω* threshold can be derived with specific considerations of the gene family at hand and the experimental conditions (e.g., the number of homologous genes, the sequence homology among the family members, and the number of sgRNAs that are collectively applied). Notwithstanding, the application of multiple sgRNAs simultaneously within a multiplex system also comes at the cost of potential efficacy reduction [53] as well as a higher number of off-targets. Moreover, increasing the frequency of potential cleavage sites enhances the chances to chromosomal translocations and may not be desired.

The CRISPys algorithm presented in this study was focused on the design of an optimal sgRNA that could best target the entire gene set. Yet, frequently a single sgRNA applicable to the entire gene family could not be found (e.g., Table 2). In such cases, a possible approach would be to partition the input gene family to subgroups according to sequence or functional similarity and to apply CRISPys on each subgroup. Nevertheless, this approach could lead to suboptimal design since gene partitioning according to homology (or functionality) does not guarantee the availability of similar targets within each partition that would provide the minimal and most efficient set of sgRNAs. Thus, in any event that a simultaneous application of multiple sgRNAs is desired, an alternative strategy would aim to design the minimal set of sgRNAs that target the entire gene set with highest efficiency. Such an approach could be translated to the set cover [54] problem, which is well studied in computer science and complexity theory. Accordingly, given a set of elements, each belongs to one or more sub-collections, the task is to identify the smallest number of sub-collections whose union covers the set. In our case, each element represents a gene. Several genes belong to a sub-collection if and only if they can be cleaved by the same sgRNA above a specified threshold. Therefore, solving the set cover problem would provide the optimal and minimal set of sgRNAs that can cleave all of the genes above a desired threshold of efficiency. Since the set cover problem cannot be solved in a polynomial time (i.e., it is NP-hard), a solution can be obtained using an approximation algorithm [55–57] or by combinatorial optimization [58]. This alternative should be useful for future genome editing applications, and its implementation is left for future research.

## Materials and Methods

### Converting the scoring function to a distance metric

In order to construct a hierarchical clustering of the target set *T*, the distance between every two targets must be computed. In the context of the CRISPR-Cas9 system, however, available scoring functions assess the cleavage propensity of a nuclear target by a given sgRNA rather than the distance between two nuclear targets. A naïve approach would be to set one of the targets as the sgRNA and compute its propensity to cleave the other target. But such an approach will not result in a valid distance metric (e.g., the scoring function is not necessarily symmetric nor does it satisfies the triangle inequality). We thus implemented two alternative procedures for converting a scoring function *φ* to an Euclidean distance function. The first alternative corresponds to a scoring function, like the CFD score [25], that treats each position independently, such that each mismatch is penalized according to the type and position of the mismatch, and the resulting score is a multiplication over all individual positions. Specifically, given a 20-nt long target *t*_*i*, a vector of length 80 is constructed in which every four entries correspond to a position in *t*_*i*; the first entry specifies the penalty according to *φ* if ‘A’ was placed in this position in the opposing sgRNA, the second stands for ‘C’, etc. Given this vector representation, the distance between two targets is calculated as the Euclidean distance between the two corresponding vectors. Thus, targets that are similar to one another receive similar scores for the different substitution possibilities in every position (dictated by *φ*), which in turn leads to a low Euclidean distance.

The second conversion was implemented to deal with any scoring function, without relying on its specific characteristics. The scoring function provided by CRISTA [37] is one example where the cleavage propensities are computed based on a non-linear combination of features. This conversation is based on transforming every target to a multidimensional space, where it is represented by a vector of cleavage propensities by a large set of sgRNAs, *S*. Specifically, for every target *t*_*i* ∈ *T*, its cleavage propensity *φ*(*s, t*_*i*) by each sgRNA *s* ∈ *S* is computed. Here, we defined *S* to be the set of sgRNAs that perfectly match the targets in *T* but other possibilities, such as a randomly-generated set of sgRNAs can be used. This produces a representative vector of length |*T*| for every target in the new space. The distance between targets *t*_*i* and *t*_*j* is calculated as the Euclidean distance between their corresponding vectors. Thus, a couple of targets that are expected to be cleaved with similar efficiencies by a set of sgRNAs will be converted to vectors with similar values, leading to a low Euclidian distance compared to targets that are expected to be cleaved dissimilarly by the same set of sgRNAs. We note that while this conversion procedure is more computationally demanding than the former technique detailed above, the two approaches yield similar results (Supplementary Text S1).

### sgRNA design for large gene families

Mutating a large number of genes from a specific family has the potential to overcome functional redundancy and to reveal the function of the encoded proteins. However, mutating a large number of genes simultaneously may lead to a lethal or sterile phenotype, limiting our ability to elucidate their function. In such cases, mutating a smaller portion of the group is desired. Dividing the input family into smaller groups according to sequence similarity would increase the flexibility of the screen and allow a more focused experimental design. To this end, we implemented a strategy that recursively splits the group of genes into homologous subgroups and generates potential sgRNAs for each of them. Specifically, a hierarchical clustering of the input gene set is constructed using the UPGMA algorithm as implemented in Biopython [59] (we note that this tree represents the similarities among the input genes, while the hierarchical clustering detailed in section “Algorithm description” represents the similarities among the targets). The input distance matrix is computed using Protdist [60] given a multiple sequence alignment generated by MAFFT [61], with its default options, on the translated genes. CRISPys is then applied to each node of the constructed UPGMA tree, producing an optimal sgRNA for each homologous subgroup.

### Program availability

An online version of the CRISPys algorithm described here is freely available at http://multicrispr.tau.ac.il/. The server accepts as input a set of (potentially homologous) sequences for which sgRNA candidate should be designed. In order to avoid targeting the designed sgRNA at intron-exon junctions, users may provide each gene as a set of exon sequences. The webserver allows users to choose between the two optimization criteria of CRISPys (*s*^*exp* and *s*^*Ω*), to choose among several available functions that determine the cleavage propensity of a DNA target by a given sgRNA (with the CFD score [25] being the default), and to optionally consider the homologous relationships among the input genes in the sgRNA design (see Methods: *‘sgRNA design for large gene families*’). Once the sgRNAs design process is completed, the website provides the possibility to search off-targets through CRISPOR [47] or CRISTA [37] web-servers.

## Supplementary Materials

**Table S1.** The effect of applying different Ω thresholds on the s^Ω^ design strategy. For each family size of 2-10 genes in the genome of *S. lycopersicum*, CRISPys was applied with the criterion. The results presented in the main text (Tables 1-2) were obtained by applying the *s^Ω^* design strategy with *Ω* = 0.43. This threshold corresponds to the 50^th^ percentile of the CFD scores over a set of experimentally validated targets (see Results, section 1.2). The table below provides the results of applying more stringent (*Ω* = 0.66) or lenient (*Ω* = 0.33) thresholds, corresponding to the 90^th^ and 25^th^ percentiles, respectively. Cleavage expectation is the average expected number of genes predicted to be cleaved across all families of that size; *c^Ω^* is the average number of genes that are predicted to be cleaved above the specified threshold (the standard deviation of each statistic is given in parentheses); percentage of fully cleaved families is the fraction of families for which an sgRNA that can cleave all family members could be successfully designed.

|  | **$s^\Omega$ designed using $\Omega = 0.66$** |  |  | **$s^\Omega$ designed using $\Omega = 0.33$** |  |  |
| --- | --- | --- | --- | --- | --- | --- |
| **Family Size** | **Cleavage expectation** | **$c^\Omega$** | **Percentage of fully cleaved gene families** | **Cleavage expectation** | **$c^\Omega$** | **Percentage of fully cleaved gene families** |
| **2** | 1.7 (0.3) | 1.68 (0.4) | 69% | 1.9(0.3) | 1.7 (0.3) | 89% |
| **3** | 2.15 (0.5) | 2.18 (0.6) | 29% | 2.55 (0.5) | 2.12 (0.5) | 56% |
| **4** | 2.56 (0.7) | 2.63 (0.7) | 15% | 3.15 (0.7) | 2.51 (0.7) | 36% |
| **5** | 2.83 (0.7) | 2.89 (0.8) | 5% | 3.71 (0.9) | 2.72 (0.8) | 23% |
| **6** | 3.16 (0.9) | 3.24 (1.0) | 3% | 4.25 (1.0) | 3.05 (1.0) | 14% |
| **7-8** | 3.64 (1.2) | 3.66 (1.3) | 3% | 3.56 (1.3) | 4.99 (1.4) | 9% |
| **9-10** | 3.92 (1.2) | 3.84 (1.3) | 0% | 3.7 (1.2) | 5.55 (1.7) | 1% |

### Text S1. Comparison of the two procedures for transforming the scoring function to Euclidean distance

To compare the two alternative procedures for converting the scoring function *φ*(*sgRNA, target*) to a distance metric between two targets (see methods, section “*Converting the scoring function to a distance metric*”), a set of 500 gene families of size 2-10 from the genome of *S. lycopersicum* was randomly sampled. For each gene family, CRISPys was executed with both procedures using the *s^exp^* design strategy. This resulted in two sgRNAs *s*^1^ and *s*^2^ per gene family, corresponding to the order of the two procedures detailed in the main text. We then compared the two designed sgRNAs using their relative difference, computed as:

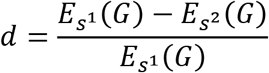

where *G* represents the gene family, and *E_s_*(*G*) represents the cleavage expectation across all genes in *G* by sgRNA *s*.

The average of the *d* values across the 500 sampled families was 0.006 with standard deviation of 0.221, and a 95% credible interval (CI) of (*−0.013, 0.025*). Since zero is contained within the CI, there is no statistical support for any difference in the results obtained by the two conversion procedures.

